# CAS12e (CASX2) CLEAVAGE OF CCR5: IMPACT OF GUIDE RNA LENGTH AND PAM SEQUENCE ON CLEAVAGE ACTIVITY

**DOI:** 10.1101/2023.01.02.522476

**Authors:** David A. Armstrong, Taylor R. Hudson, Christine A. Hodge, Thomas H. Hampton, Alexandra L. Howell, Matthew S. Hayden

**Author notes:** Corresponding authors: Alexandra L. Howell, PhD., VA Medical Center, 215 North Main St, White River Jct., VT 05009, Email address; and Matthew S. Hayden, M.D., Ph.D., Department of Dermatology, Dartmouth Health, Lebanon, NH 03756. Email addresses: David A. Armstrong; Taylor R. Hudson; Christine A. Hodge; Thomas H. Hampton; Alexandra L. Howell; Matthew S. Hayden.

## Abstract

CRISPR/Cas is under development as a therapeutic tool for the cleavage, excision, and/or modification of genes in eukaryotic cells. While much effort has focused on CRISPR/Cas from *Streptococcus pyogenes* (SpCas9) and *Staphylococcus aureus* (SaCas9), alternative CRISPR systems have been identified using metagenomic datasets from non-pathogenic microbes, including previously unknown class 2 systems, adding to a diverse toolbox of gene editors. The Cas12e (CasX1, CasX2) endonucleases from non-pathogenic Deltaproteobacteria (DpeCas12e) and Planctomycetes (PlmCas12e) are more compact than SpCas9, have a more selective protospacer adjacent motif (PAM) requirement, and deliver a staggered cleavage cut with 5-7 base overhangs. We investigated varying guide RNA (spacer) lengths and alternative PAM sequences to determine optimal conditions for PlmCas12e cleavage of the cellular gene *CCR5* (CC-Chemokine receptor-5). *CCR5* encodes one of two chemokine coreceptors required by HIV-1 to infect target cells, and a mutation of *CCR5* (delta-32) is responsible for HIV-1 resistance and reported cures following bone marrow transplantation. Consequently, *CCR5* has been an important target for gene editing utilizing CRISPR, TALENs, and ZFNs. We determined that *CCR5* cleavage activity varied with the target site, guide RNA length, and the terminal nucleotide in the PAM sequence. Our analyses demonstrated a PlmCas12e PAM preference for purines (A, G) over pyrimidines (T, C) in the fourth position of the CasX2 PAM (TTCN). These analyses have contributed to a better understanding of CasX2 cleavage requirements and will position us more favorably to develop a therapeutic that creates the delta-32 mutation in the *CCR5* gene in hematopoietic stem cells.

## INTRODUCTION

The C-C chemokine receptor 5 (CCR5) was identified as a critical co-receptor for target cell infection by macrophage-tropic primary isolates of HIV-1 [1, 2]. Since then, the *CCR5* gene or its protein receptor, have been targets for modification as an approach to inhibit or prevent HIV-1 infection of T lymphocytes and myeloid cells [3]. Early attempts to mutate or delete *CCR5* used TALENs (transcription activator-like effector nucleases) [4], or ZFNs (zinc finger nucleases) [5], as approaches to excise critical gene regions within the *CCR5* coding segments [6-11]. Another approach to modify *CCR5* gene expression has been reported using base editors [12, 13], a technique in which altering the sequence of nucleotides in the gene sequence can lead to the insertion of targeted point mutations or to the introduction of premature stop codons.

More recently, genome editing using CRISPR/Cas has been used to achieve precise gene cleavage of both cellular and integrated viral genes in eukaryotic cells and in pre-clinical animal models [14-19]. This method provides a powerful approach to eliminate, modify, or replace specific genes or gene sequences, using a combination of a Cas (CRISPR-associated) endonuclease, and a short RNA molecule known as the guide RNA (gRNA, or “spacer”). For example, CRISPR/Cas gene editing has provided novel approaches for the development of therapeutics to treat sickle cell disease (CTX001), HIV-1 (EBT-101), and non-small cell lung cancer [20], with several other clinical trials in the pipeline. The key to CRISPR’s specificity lies in the binding of the gRNA to its complementary target gene sequence adjacent to the protospacer adjacent motif (PAM) that serves to license the Cas endonuclease activity. Target site selection is based on multiple parameters. For example, the presence of a PAM dictates the location and orientation of potential target sites. The DNA strand that contains the PAM is referred to as the non-target strand (NTS), and the guide RNA binds to the DNA strand without the PAM known as the target strand (TS). Further, the similarity of the target sequence to other genomic sites must also be considered to reduce the potential for off-target cleavage events. *CCR5* targeting is challenging in this regard, given the homology of *CCR5* to *CCR2* [21], another C-C Chemokine receptor. CRISPR/Cas cleavage of *CCR5* in hematopoietic cells was shown to cause loss of receptor expression on differentiated progeny cells [22]. Targeting *CCR5* in hematopoietic stem cells as an approach to generate a CCR5^null^ immune system was supported by clinical reports of permanent suppression of HIV replication in two different patients who underwent allogeneic bone marrow transplantation using bone marrow from donors homozygous for the *CCR5*^*Δ32*^ mutation [23-26]. To date, the most common sources of Cas endonucleases, Cas9, derive from the common human pathogens *Staphylococcus aureus* (SaCas9) and *Streptococcus pyogenes* (SpCas9), yet pre-existing immunity to these bacteria as well as the rather large molecular size of SpCas9 may limit the long-term utility of these editors in therapeutic approaches [27-29].

Recent metagenomic sequencing and computational efforts have identified a wide variety of new CRISPR systems, and have led to the discovery of Cas12a [30, 31], Cas12b [32, 33], Cas12d [34, 35], Cas12e [35, 36], and the RNA-cleaving Cas13 family [37]. Cas12e (DpbCasX1 and PlmCasX2; formerly referred to as CasX1 and CasX2, respectively), were identified along with Cas12d (CasY) from a metagenomic survey by Burstein, et al., [35]. Further analyses of Cas12e editors demonstrated efficient RNA-guided cleavage using a 20 nucleotide (nt) crispr RNA (crRNA) complexed to a tracrRNA [38]. Cleavage of the DNA target occurred adjacent to a PAM sequence of TTCN, and cleavage products were identified on both the target and non-target strands in an asymmetrical (staggered) pattern characteristic of Cas enzymes with a single RuvC cleavage site [39].

We sought to define optimal conditions for Cas12e (PlmCasX2) cleavage of DNA, using as a target, a 1,250 base pair (bp) fragment of *CCR5* [40] flanking the area that includes the region deleted in the delta-32 mutation [41]. Our interest in optimizing CasX2 cleavage stemmed from the potential to use this editor to replace the relevant wild-type region of *CCR5* with the delta-32 mutation. Several groups have investigated CasX cleavage activity [36, 39, 42] since its initial description by Burstein et al., [35]. Liu, et al., [39] tested cleavage with CasX1 (DpbCasX) and CasX2 (PlmCasX) and determined that a 20 nt guide RNA cleaved 12-14 nt after the PAM on the non-target strand, and 22-25 nt after the PAM on the target strand, generating a 7-14 nt 5’ overhang. They observed no single-stranded nuclease activity, unlike what is reported for Cas12a. Subsequently, Selkova, et al., [42] tested the locations of the DpbCasX staggered cleavage cuts using guide RNAs of various lengths and high throughput sequencing. Contrary to findings by Liu, et al., [39], this group observed cleavage of the non-target strand at 17-19 nts from the PAM, and after the 22 nt on the target strand, producing shorter 5’ overhangs of only 3-5 nt lengths. However, they also observed that shortening the guide RNA from 20 nt to 16 nt shifted the cleavage site on the non-target strand closer to the PAM, producing longer 5’ overhangs of 6-8 nt in length. In a recent study by Tsuchida, et al., [36] homologs of CasX and single guide (sg) RNAs were found to impact cleavage efficiency of these editors. This group observed enhanced *in vitro* cleavage of target with DpbCasX compared to PlmCasX, but improved cleavage with PlmCasX in cells suggesting *in vitro* cleavage activity is impacted by the purity of the enzyme prep. Further, they determined that three nucleotide-binding loops within CasX may enhance PAM-proximal region recognition, sgRNA interaction, and DNA substrate loading, all of which may contribute to the different cleavage efficiencies among homologs of CasX [36].

To further understand factors influencing CasX2 function, we performed a systematic analysis of *CCR5* gene cleavage by multiple guide RNAs of varying lengths. We also analyzed the impact of changes in the terminal PAM base, (the “N” in TTCN), to determine the extent to which CasX2 exhibits any base preference at this site. Our findings showed that cleavage efficiencies varied with both the guide RNA target site, the length of the guide RNA, and the terminal PAM base. These findings underscore the need for a more systematic analysis of CasX2 target selection and gRNA design in order to improve therapeutic implementation of these new CRISPR enzymes.

## METHODS

### Recombinant CasX2 protein expression and purification

CasX2 expression plasmids were generated by ligation independent cloning. The CasX2 ORF was amplified from pBLO 62.5 (pBLO 62.5 was a gift from Jennifer Doudna and Benjamin Oakes, Addgene plasmid #123124; http://n2t.net/addgene:123124; RRID:Addgene 123124) [39]. The amplicons containing CasX2 were subcloned into the pET His10 MBP Asn10 TEV LIC cloning vector (pET His10 MBP Asn10 TEV LIC cloning vector (2CT-10) was a gift from Scott Gradia, Addgene plasmid #55209; http://n2t.net/addgene:55209; RRID: Addgene_ 55209), and verified through Sanger sequencing. The final plasmid expressed CasX2 with a TEV-cleavable N-terminal His10-MBP fusion tag.

His10-MBP-CasX2-2CT-10 was transformed and expressed in Rosetta™(DE3) Competent Cells (Novagen, St. Louis, MO). Cells were grown at 37°C in 1 liter LB broth supplemented with ampicillin (100 μg/ml), chloramphenicol (34μg/mL), and 0.1% glucose. After growing to an OD_600_ of 0.5, the temperature was decreased to 16°C, and CasX2 expression induced with the addition of 0.5 mM isopropyl β-D-1-thiogalactopyranoside (IPTG, EMD Millipore, Burlington, MA). After 16 hours at 16°C, bacterial cells were pelleted and resuspended in lysis buffer (50mM HEPES, pH 7.5, 0.5 M NaCl, 0.5 mM tris-(2-carboxyethyl)-phosphine (TCEP), 20 mM imidazole, 10% (v/v) glycerol, 1 mM phenylmethyl-sulfonyl fluoride (PMSF), 1 tablet protease inhibitor cocktail (Sigma Aldrich, St. Louis, MO) per 50 mL of solution, and 0.5 mg/mL lysozyme), at a volume of 5 mL lysis buffer per gram of pellet. The resuspended cells were disrupted by sonication, and cellular debris removed by ultra-centrifugation for 30 minutes at 35,000 x g. The supernatant was loaded onto a 5 mL HisTrap HP cartridge (#17524802, Cytiva Life Sciences, Marlborough, MA) using an AKTA Go Chromatography system (#29383015, Cytiva Life Sciences). Columns were washed with 50 mL Ni Wash Buffer 1 (50mM HEPES, pH 7.5, 0.5 M NaCl, 0.5 mM tris (2-carboxyethyl) phosphine (TCEP), 20 mM imidazole, 10% (v/v) glycerol) and 50mL Ni Wash Buffer 2 (50mM HEPES, pH 7.5, 0.6 M NaCl, 0.5 mM TCEP, 30 mM imidazole, 10% (v/v) glycerol), and eluted in 1.5 mL fractions with 15mL Ni Elution Buffer (50mM HEPES, pH 7.5, 0.5 M NaCl, 0.5 mM TCEP, 400 mM imidazole, 10% (v/v) glycerol). Fractions containing the peak elution products (as determined by spectrophotometry) were combined, and diluted in 2x MBP Wash Buffer (50mM HEPES, pH 7.5, 0.5 M NaCl, 0.5 mM TCEP, 10% (v/v) glycerol). To further purify CasX2, the diluted Ni column elution was subjected to a gravity fed column packed with 4 mL Amylose Resin (# E8021, New England Biological, Ipswich, MA), washed with 50mL of MBP wash buffer, and eluted in 1mL fractions with MBP Elution Buffer (50mM HEPES, pH 7.5, 0.5 M NaCl, 0.5 mM TCEP, 10% (v/v) glycerol, and 20 mM maltose).The eluted samples were visualized via SDS-PAGE and fractions containing the target product were combined and dialyzed overnight at 4°C against storage buffer (50mM HEPES, pH 7.5, 0.5 M KCl, 0.5 mM TCEP, and 20% (v/v) glycerol). The protein product was recovered, concentrated using 50 kDa MWCO Amicon Ultra Centrifugation Filter units (Millipore, Burlington, MA), and stored in aliquots at -80°C.

### Design of guide RNAs

Human *CCR5* (NM_000579.4), located on chromosome 3, was selected as the target sequence for CasX2 cleavage. Single guide RNAs (sgRNA) were designed using Benchling (www.benchling.com) to search for target regions adjacent to a 5’ TTCN PAM sequence. Guide RNA sequences were individually selected for high specificity scores, with an off target cut off of >92.0, and proximity to the wild-type delta-32 sequence within exon 2 of *CCR5* (Chr3:46372947-46373940). We selected five target sequences upstream and five target sequences downstream of the delta-32 mutation site on *CCR5* as regions for guide RNA binding.

### *In vitro* T7 sgRNA transcription

Transcription templates for sgRNAs were constructed with a 5’ T7 promoter and the CasX2 scaffold sequence followed by the selected 17-23 bp sequence complementary to the target DNA sequence. The template was generated through overlap extension PCR using primers that were purchased from Integrated DNA Technologies (IDT, Coralville, IA). An initial annealing step was performed with a 110 nt forward primer, X2_F_template (5’-GAATTAATACGACTCACTATAGTACTGGCGCTTTTATCTCATTACTTTGAGAGCCATCACCAGCGACTATGTCGTATGGGTAA AGCGCT-3’) containing the sequences for the T7 promoter and tracr region, and a reverse primer, X2_R_template (5’-(17-23 bp gRNA) + CTTTGATGCTTCTTATTTATCGGATTTC TCTCCGATAAATA-3’) containing the variable gRNA, by combining 45 μM of the forward and reverse primers in 1X STE buffer for a total volume of 10 μl, and annealing for 5’ at 95°C followed by slow cooling to room temperature before the addition of 90 μl of nuclease free-H_2_O. Post-annealing PCR was performed with the addition of a shorter forward primer (5’-GAAATTAATACGACTCA CTATAGTACTGGCGCTTTTATCT-3’) and the X2_R_ Template in 10 μM final concentration with 5 μl polymerase master mix at the following cycling conditions: 98°C/30sec, 25 x (98°C/5sec, 68°C/10sec, 72°C/15sec), 72°C/2min, 4°C. The PCR product was purified with the Qia-quick PCR clean up kit (Qiagen, Germantown, MD). The Hi-Scribe T7 High Yield RNA synthesis kit (NEB) was used to generate RNA transcripts at 37°C for 16 hours, according to the manufacturer’s protocol. The final sgRNA sequence is: 5’**-**UACUGGCGCUUUUAUCUCAUUACUUUGAGAG CCAUCACCAGCGACUAUGUCGUAUGGGUAAAGCGCUUAUUUAUCGGAGAGAAAUCCGAU AAAUAAGAAGAUCAAAG + (17-23 nt gRNA)-3’. Sequences of the DNA oligonucleotides used for *in vitro* transcription, as well as the RNA sequences of the resulting 17-23 bp spacers of the sgRNAs are shown in Supplemental Table 1.

### DNA target generation for *in vitro* cleavage (IVC) reactions

The 6,415 bp pcDNA3-CCR5 plasmid was a gift from Erik Procko (Addgene plasmid #98943; http://n2t/net/addgene:98943; RRID:Addgene_98943) [43]. Prior to use in IVC, a smaller target fragment of 2,812 bp was generated by sequential restriction digests with SmaI/ SpeI, and subsequent purification via Qia-Quick PCR clean up columns (Qiagen), according to the manufacturer’s protocol. The restriction digested DNA was purified and quantitated using a Qubit dsDNA assay.

### *In vitro* cleavage

*In vitro* cleavage (IVC) reactions were performed in a two-step process. First, 10 μl of a ribonucleoprotein complex (RNP) was formed by incubating sgRNA and CasX2 at equimolar ratios in 1x IVC buffer (1x IVC buffer: 20mM HEPES pH 7, 100mM KCl, 5mM MgCl_2_, 0.1mM EDTA, 1% glycerol) at 37°C for 10 mins. After RNP formation, the DNA target was added in an 80:1 RNP:DNA ratio (unless otherwise specified), in a 20 μl volume, and incubated at 37°C for 60 min. Various RNP:DNA molar ratios, from 10:1 to 80:1, were chosen for each RNP to achieve cleavage in a linear range. Agarose gel (1%) electrophoresis was used to visualize IVC products. Gels were visualized using a Bio-Rad ChemiDoc MP imaging system and the percent cleavage of the target sequence determined via densitometry analysis using Bio-Rad Imaging Lab software (V6.1.0 Build7-2020).

### Design of IVC DNA target sequences with altered PAM terminal bases

Four separate gene blocks (gBlocks) corresponding to a 1,114 bp region of the human *CCR5* gene sequence that included nine of the ten sgRNA target sequences, were synthesized by IDT. For these fragments, the terminal PAM base of each target for nine of the sgRNA sites was changed to either an adenine (A), cytosine (C), guanine (G) or thymidine (T), resulting in four target sequences with the only difference being the alternate terminal PAM base substitutions. Each of the four gene fragments were cloned into the plasmid pcDNA3. Briefly, the fragments and plasmid were treated with the NheI and XhoI restriction enzymes and ligated with T4 DNA ligase in a 3:1 molar ratio before transforming into NEB® 5-alpha competent *E. coli*. Selected colonies were screened and verified by Sanger Sequencing.

## RESULTS

### *In vitro* cleavage efficiency of single guide RNAs (sgRNA) targeting *CCR5*

Ten different sgRNAs with specificity to exon 2 of *CCR5* (Chr3:46372947-46373940) were selected to assess *in vitro* cleavage activity of PlmCas12e (Table 1). SgRNAs 1, 2, 4, 7 and 10 bound to the (-) target DNA strand, while guides 3, 5, 6, 8 and 9 bound to the (+) target DNA strand. Specificity scores, as determined in Benchling.com and based on Hsu et al., [44], ranged from 92.825 (sg 2) to 98.825 (sg 9). Three sgRNAs used a PAM sequence of TTCA, three used a PAM sequence of TTCC, and four used a PAM sequence of TTCG. Single guide RNAs were selected based on their location relative to the CCR5 Δ32 site, with five sgRNAs upstream of the Δ32 site, and five sgRNAs downstream of the Δ32 sequence as depicted in Figure 1. Additional sequence information and location of the guides relative to each other and to the CCR5 Δ32 sequence is shown in Supplemental Figure 1.

**Table 1.** CCR5 sgRNA Position on Chromosome 3 and Specificity Score

| RNA<br>Guide | Position | Strand | Specificity<br>Score | PAM |
| --- | --- | --- | --- | --- |
| 1 | 46372957 | (-) | 93.552 | TTCA |
| 2 | 46373065 | (-) | 92.825 | TTCA |
| 3 | 46373164 | (+) | 93.068 | TTCT |
| 4 | 46373172 | (-) | 98.547 | TTCT |
| 5 | 46373178 | (+) | 98.657 | TTCC |
| 6 | 46373664 | (+) | 97.692 | TTCT |
| 7 | 46373680 | (-) | 95.179 | TTCA |
| 8 | 46373897 | (+) | 96.788 | TTCT |
| 9 | 46373904 | (+) | 98.825 | TTCC |
| 10 | 46373921 | (-) | 96.577 | TTCC |

**Figure 1.**
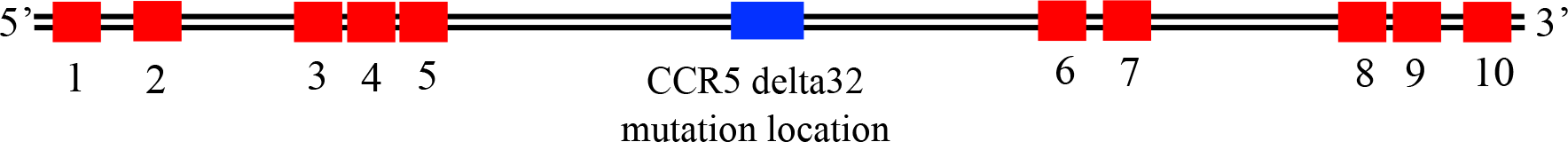
Target sequences for sgRNAs within the human *CCR5* gene. Schematic of the relative locations of the target sequences for sgRNAs (red rectangles) relative to the location of the region that would be deleted in the Δ-32 mutation (blue), within a 1,000 base pair segment of exon 2 of the human *CCR5* gene located on chromosome 3.

A previous report by Selkova, et al., [42] investigated the correlation between spacer length and location of the cleavage cuts on DNA targets. In their studies, they noticed slight differences in the efficiency of cleavage of one target as the spacer lengths were shortened. Based on their observation, we sought to carry out a systematic evaluation of spacer length on cleavage efficiency of ten different target sites to determine if a correlation existed between these variables. We used a 2,812 bp (Sma I/Spe I) fragment of *CCR5* derived from the plasmid pcDNA3-CCR5 that included all ten target sites. *In vitro* cleavage reactions of the *CCR5* target were performed for all ten sgRNAs at spacer lengths from 17-23 nt, and each experiment was repeated for a total of three times with consistent findings. IVC reactions were then subjected to agarose gel electrophoresis, where cleavage of the target could be quantified.

Figure 2A shows a representative experiment for sg 7, and the results for the other nine sgRNAs are shown in Supplemental Figure 2. We found that cleavage efficiencies varied with both the sgRNA target as well as the length of the sgRNA. Although the spacer length had a significant impact on the efficiency of some sgRNAs, we were unable to identify a consistent correlation between spacer length and cleavage activity across the 10 sgRNAs we tested. For example, sg 3 cut most efficiently at spacer lengths 18, 19 and 23 nt, sg 7 cut most efficiently at lengths from 17-20 nt, and sg 9 cut most efficiently at a spacer length of 18 nt.

**Figure 2.**
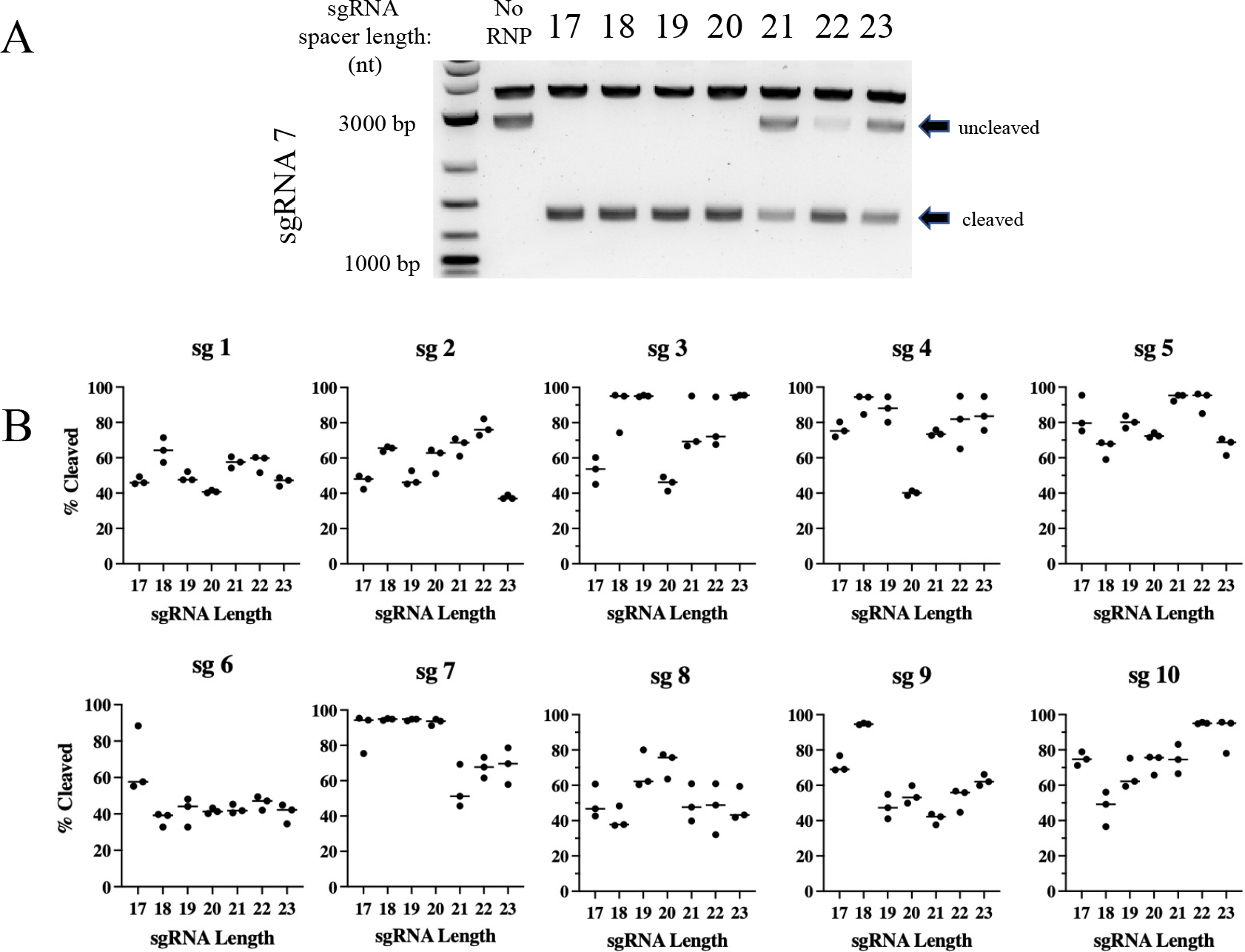
Cleavage activity of Cas12e is spacer length and target location dependent. **A)** Agarose gel separation was used to visualize Cas12e cleavage products after *in vitro* cleavage. A *CCR5* gene target sequence of 2,812 nucleotides (nt) was used to examine the effect of spacer length on DNA cleavage. A representative gel for sgRNA 7 is depicted as an example from which densitometry measurements were derived. The sgRNA 7 generated cleavage products of 1,368 nt and 1,444 nt in length. **B)** Seven different spacer lengths from 17 nt to 23 nt for each of ten different sgRNAs were analyzed for *in* vitro cleavage activity of the 2,812 nt *CCR5* DNA target by agarose gel electrophoresis. The densitometry of the cleavage products was analyzed for each sgRNA at each spacer length, and the median percent cleavage of the target plotted in a dot blot using *Prism Soft Inc*. All experiments were run in triplicate.

Densitometry analysis was performed to determine the cleavage activity of each guide RNA at all lengths examined. The median percent target sequence cleaved was plotted against sgRNA length from all sgRNAs and shown in Figure 2B. Characteristics of each sgRNA were noted, including sgRNA length, GC content, PAM sequence, cutting efficiency and Gibb’s free energy (δG) (Supplemental Table 2). We assessed cleavage activity as a function of spacer length to determine whether some spacer lengths were significantly more efficient than others. A two-way ANOVA of cleavage activity as a function of spacer length and guide RNA did not reveal significant differences in cleavage efficacy when p values were adjusted for multiple comparisons using Tukey’s honest significant differences; all p values exceeded 0.05.

### PAM sequences impact sgRNA cleavage activity

Given that sgRNA length impacted cleavage activity *in vitro*, we sought to determine whether the PAM sequence could also influence the performance of the sgRNA. Considering that the PAM for CasX2 is TTCN, the terminal base of the PAM can be any nucleotide. For these experiments, we synthesized four different target DNA sequences in which the 4^th^ nucleotide of the PAM for each sgRNA target was either an A, G, C or T nucleotide. Figure 3 depicts the location of each of the modified bases in the target, where “N” represents the terminal PAM for each of the sgRNAs. Four substitutions were located upstream of the *CCR5* Δ32 sequence, and five substitutions were located downstream. One sgRNA was not evaluated (sg 3) because the PAM change for sg 4 was located within the sg 3 binding region, and therefore created a mismatch between sg 3 and its target. We chose several sgRNAs at varying lengths to assess cleavage of each of the four DNA targets. We observed that for each sgRNA, cleavage activity varied dramatically according on the terminal PAM base in the target DNA. For example, sg 7 at a length of 18 nt cleaved target DNA more efficiency when the PAM sequence ended with either an A or G, compared to the same target when the terminal PAM base was a C or a T (Figure 4A). This finding was consistent for many of the sgRNAs regardless of spacer lengths tested, and indicated that a purine in the terminal position of the PAM was generally preferred for target cleavage. Normalized percent cleavage from six different biological replicates is shown in Figure 4B. Additional replicates with other sgRNA lengths are shown in Supplemental Figure 3. When we examined the differences in Cas12e cleavage comparing the terminal PAM base via linear modeling, the analysis revealed for percent cleavage as a function of sgRNA, spacer length, and terminal PAM nucleotide base, both C and T are significantly less effective than G (e.g., C (p < 0.0031), and T (p < 0.00012)).

**Figure 3.**
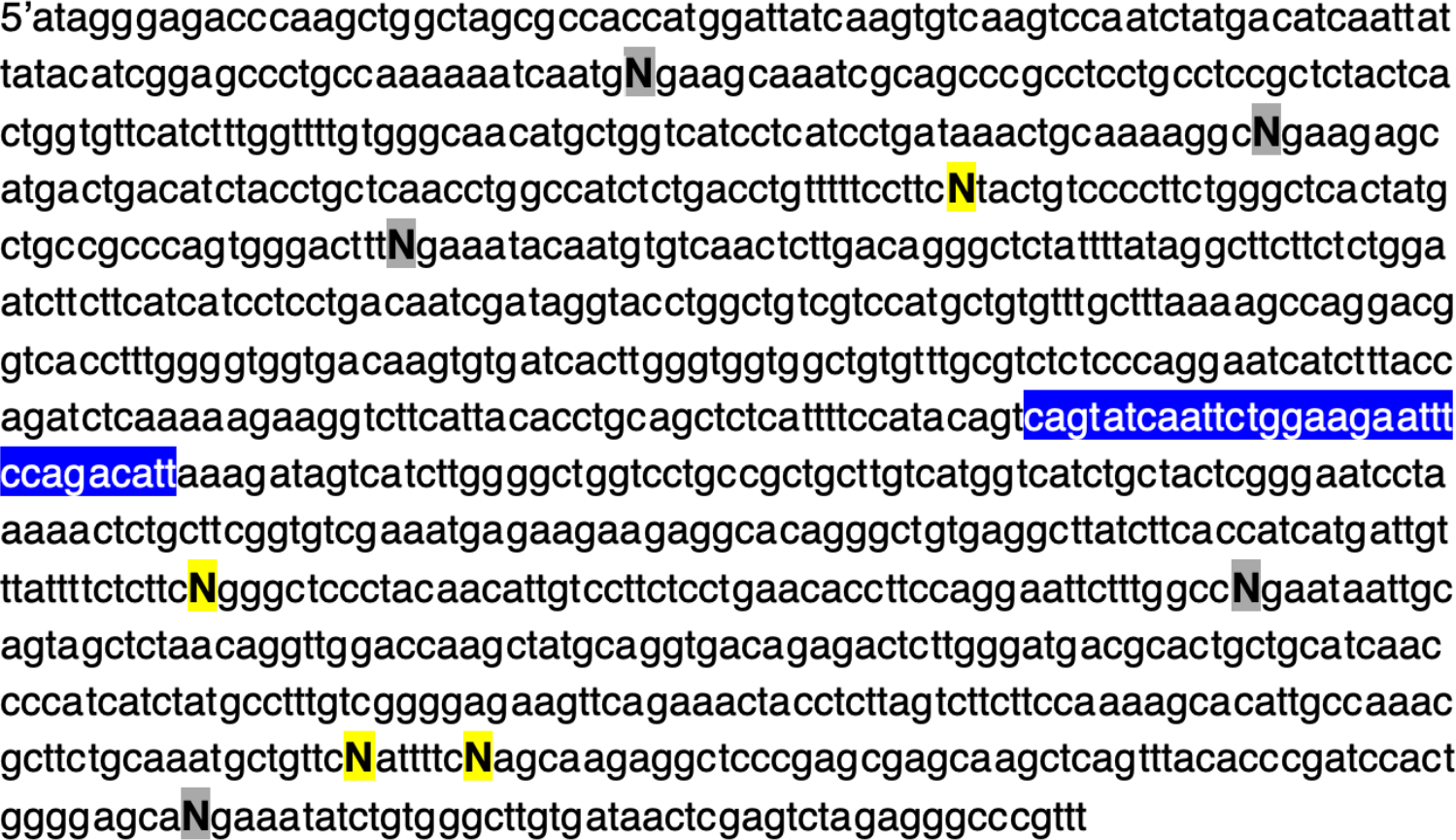
Location of nine terminal PAM bases that were changed to assess CasX2 PAM preference. Four different *CCR5* gene fragments of 1,114 bp (gBlocks) each were synthesized to include nine of the ten gRNA target regions. The terminal PAM base for each sgRNA was changed to either an A, C, G or T in each gBlock. Each of the four gene fragments were separately cloned into the pcDNA3.1 vector and then restriction enzyme digested with NheI and XhoI to yield a 1,074 bp target. A yellow highlighted N (**N)** indicates a terminal base on the (+) strand, and a grey highlighted N (**N**) indicates a terminal base on the (-) strand. The blue highlighted region is the wild-type sequence of *CCR5* that would be deleted in the Δ-32 mutation. The sgRNAs 1, 2, 3 and 5 bind upstream of the Δ-32 region and sgRNAs 6, 7, 8, 9 and 10 bind downstream.

**Figure 4.**
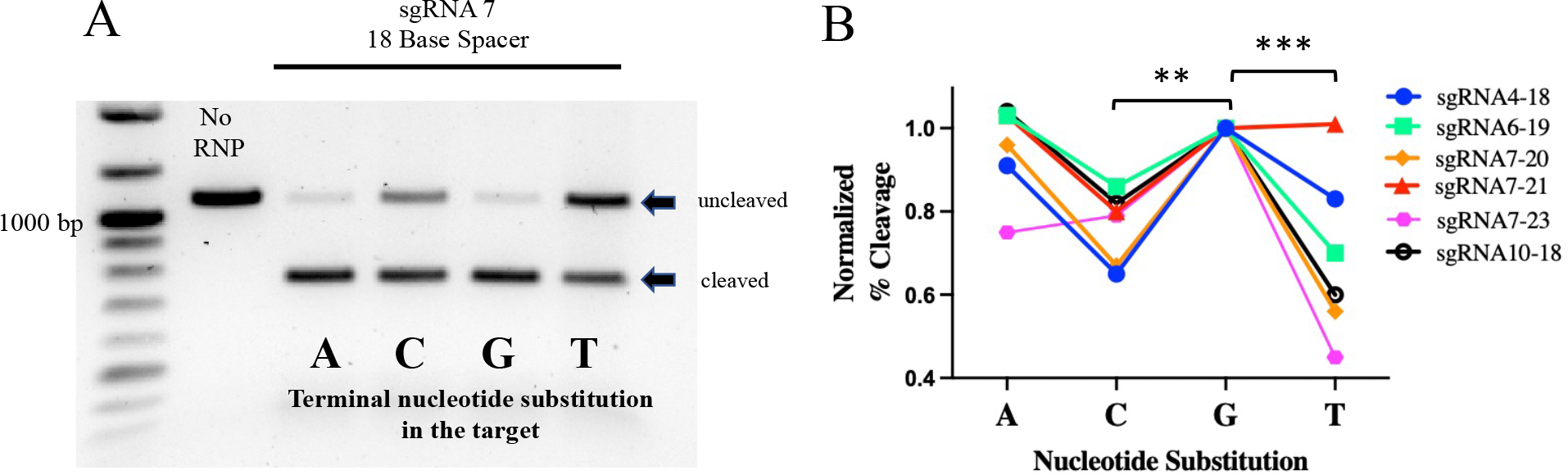
The terminal PAM base influences Cas12e cleavage activity. **A)** *CCR5* gene fragments (1,074 nt) were generated with different terminal PAM bases of either A, C, G or T. These four DNA targets were assessed for cleavage activity by sgRNA 7 at a spacer length of 18 nt. Cleavage products were then run on a 1% agarose gel so that differences in the cleavage activity for each target could be quantified. A representative experiment is shown in panel A. In triplicate experiments, cleavage activities for sgRNA 7 at spacer length of 18 nt, were highest for the terminal “G” (median cleavage 88.5%), followed by the terminal “A” (66.3%), terminal “C” (59.4%), and terminal “T” (43.6%). **B)** Cleavage activity for each terminal PAM base substitution was determined by gel densitometry and normalized to the “G” target. Results from four different sgRNAs across multiple spacer sizes shows purines A and G as the terminal PAM base demonstrated greater cleavage activity compared to C and T. Linear modeling reveals that for percent cleavage as a function of guide, guide length and base, both C and T are significantly less than G (**p < 0.01, ***p < 0.001).

## DISCUSSION

We sought to identify parameters associated with increased efficiency of PlmCas12e cleavage of DNA targets that would aid in design and development of future gRNAs for Cas12e applications. The parameters we tested included target location, gRNA spacer length, and terminal PAM base. *In vitro* cleavage reactions were used to examine cleavage activity. Our results demonstrated that both gRNA spacer length and the PAM terminal base are important factors to consider when designing Cas12e guide RNAs for gene editing.

The CRISPR/CasX2 (PlmCas12e) system was first identified from metagenomic sequencing of DNA extracted directly from natural microbial communities [35]. This CRISPR system, like many other previously characterized CRISPR systems, uses a gRNA and tracrRNA complex to bind to the Cas enzyme. This ribonucleoprotein complex then cleaves a DNA target dictated both by the complementarity of the gRNA to the target and recognition by the Cas enzyme of the PAM sequence. The CasX PAM, TTCN, was initially characterized using a single gRNA and did not resolve a strong preference at the fourth position [35]. However, a careful evaluation of the data obtained by Burstein, et al., in their *E. coli* PAM depletion assay using the highest depletion value threshold reported, does indicate a slight preference for a purine at the fourth position for PlmCas12e [35]. Our results suggest that further consideration should be given to the PAM sequences of Cas12e, and that PAM preferences should be assessed using multiple spacer/protospacer pairs in order to fully understand parameters influencing cleavage efficiency.

In addition to the PAM sequence, it was clear that other factors were likely to influence cleavage efficiency. Unlike the well-studied Cas9 systems, CasX2 delivers a cleavage cut on the target DNA that is asymmetrical. Using a 20 nt long gRNA, Liu et al., [39] reported that cleavage of the target DNA (termed the “protospacer”) occurred 12-14 nt from the PAM on the non-target strand. The target strand cut occurred several nucleotides after the gRNA binding site at position PAM+22-25. This type of asymmetrical cleavage cut results in a segment of single-stranded DNA termed a “5’ overhang”. In contrast, a subsequent publication by Selkova, et al., [42], reported that the cleavage sites produced by Cas12e on each strand of DNA were at locations different than those reported by Liu, et al., [39] and were 16-19 nts from the PAM on the non-target strand, and nearly exclusively at 22 nt from the PAM on the target strand when using a 20 nt-long gRNA. Selkova, et al., also reported that shortening the length of the gRNA altered the location of only the non-target cleavage site which moved toward the PAM, with the result being that shorter gRNAs resulted in larger 5’-overhangs [42]. Selkova, et al., did not find that sgRNA length had a significant or consistent effect on cleavage *in vitro* for the two targets they analyzed [42], although they did observe a slight diminution in cleavage efficiency as the spacer length was shortened for one of their targets. Their data did suggest, however, that the length of the guide RNA played a role in dictating the precise cleavage sites in the target region, a finding that has important consequences for guide RNA design.

We undertook a systematic analysis of the impact of guide RNA length on cleavage activity, using as a target, a portion of *CCR5* which included the area deleted in the delta-32 mutation. Ten different gRNAs were designed, with five upstream of the delta-32 CCR5 region, and five downstream. Five of the gRNAs bound to the positive DNA strand, whereas the other five bound to the negative DNA strand. Each guide RNA was transcribed at varying lengths from 17 nt to 23 nt and used in *in vitro* reactions where cleavage activity could be experimentally measured. Our findings demonstrated that cleavage activity varied depending on gRNA length for each particular gRNA, but there was no consistent pattern among the different gRNAs. Some of the gRNAs, including sg 1, sg 5 and sg 6, expressed consistent levels of cleavage regardless of gRNA length. Others, including sg 2 and sg 10, showed increased cleavage efficiency as gRNA length increased, whereas sg 7 and sg 8 showed declining cleavage efficiencies as guide length increased. We observed remarkable consistency with repeated experiments, suggesting that these findings are particular for the gRNAs we tested. The wide range of cleavage activity for particular guide RNAs observed as spacer lengths changed also indicated that factors other than gRNA binding location, or location on the (+) or (-) DNA strand, were impacting Cas12e cleavage events. Our findings clearly indicate that careful gRNA design and testing of a wide range of spacer lengths is important for selecting optimal guides for DNA cleavage.

We next assessed whether the fourth nucleotide in the PAM could affect cleavage activity of the guide RNAs. As CasX recognizes the sequence TTCN, it should cleave DNA targets adjacent to a PAM of TTCA, TTCG, TTCT and TTCC. To determine the contribution of the PAM sequence, we designed four different gBlock targets, substituting the last nucleotide in each PAM region with either an A, G, T or C. For these experiments, we chose several of the gRNAs at varying lengths. Our results showed that, in general, PAM sequences that ended in a purine, (e.g., TTCA and TTCG), demonstrated enhanced cleavage over PAM sequences ending in a pyrimidine, (e.g., TTCT and TTCC). This finding suggests that in addition to the gRNA sequence and the gRNA length, the sequence of bases that comprise the PAM also factor into cleavage efficiency. Because cleavage mediated by CasX2 is distal to the PAM, it is likely that the purine-purine bond adjacent to the gRNA binding site is not directly implicated in cleavage by the RuvC domain of CasX2, but to other, as of yet, undefined factors.

The studies reported in this manuscript addressed the optimal cleavage conditions for the novel Cas12e editor, CasX2. We chose to use the CCR5 gene as a target because of the importance of this receptor in HIV-1 infection. Our goal is to design guide pairs that flank the area of the CCR5 gene that has the delta-32 mutation in such a way that we can insert a donor fragment of DNA expressing the delta-32 mutation to replace the wild-type sequence. Future studies should focus on understanding the applicability of these findings to CasX2 cleavage activity and specificity in cells. Understanding PAM preferences and optimal guide lengths will assist us in designing a therapeutic approach with a higher probability of success.

## Data availability

The data that support the findings of this study are included in the manuscript.

## FIGURE LEGENDS

**Supplemental Figure 1. The 1,000 bp region of the *CCR5* gene containing all sgRNA target sequences.** Complete sequence of the *CCR5* gene region (Chr3:46372947-46373940) within exon 2 on chromosome 3 displaying the location of each sgRNA relative to the location that would be deleted in the Δ-32 mutation (blue rectangle). sgRNAs are shown in red and the protospacer adjacent motifs (PAM) in gray.

**Supplemental Figure 2. Cleavage activity by Cas12e (CasX2) is gRNA spacer length and DNA target location dependent**. Examples of agarose gel separation of CasX2 cleavage products by nine different sgRNAs are shown (sg2, sg3, sg4, sg6, sg9, sg1, sg5, sg8, and sg10). The number and size of the cleavage products are associated with their relative location within the 2,812 nt target region. Shown is one representative experiment for each gRNA at all spacer lengths. All experiments were run in triplicate.

**Supplemental Figure 3. Terminal PAM base impacts Cas12e cleavage activity for multiple sgRNAs**. *CCR5* DNA targets were generated to contain either an A, G, C or T in the terminal PAM location for each of the sgRNAs. Targets underwent *in vitro* cleavage to assess the impact of altering the terminal PAM base. A consistent cleavage pattern among several gRNAs showed the highest cleavage activity with an A or G as the terminal PAM base. To aid in visualizing cleavage products, different sized *CCR5* targets were utilized dependent on the guide location. For sgRNA 3 (panel A), a SpeI/XhoI 1,720 bp target was used, for sgRNA 4 (panel B) and sgRNA 7 (panel C), a NheI/XhoI 1,074 bp target was used, and for sgRNA 10 (panel D), a NheI/SmaI 2,166 bp target was used. Arrows indicate the location of the uncleaved target.

## REFERENCES

1. Dragic, T., V. Litwin, G.P. Allaway, S.R. Martin, Y. Huang, K.A. Nagashima, C. Cayanan, P.J. Maddon, R.A. Koup, J.P. Moore and W.A. Paxton, HIV-1 entry into CD4+ cells is mediated by the chemokine receptor CC-CKR-5. Nature, 1996. 381(6584): p. 667–73.

2. Deng, H., R. Liu, W. Ellmeier, S. Choe, M. Unutmaz, M. Burkhart, P. DiMarzio, S. Marmon, R. Sutton, C. Hill, C. Davis, S. Peiper, T. Schall, D. Littman and N. Landau, Identification of a major co-receptor for primary isolates of HIV-1. Nature, 1996. 381(6584): p. 661–6.

3. Barmania, F. and M.S. Pepper, C-C chemokine receptor type five (CCR5): An emerging target for the control of HIV infection. Appl Transl Genom, 2013. 2: p. 3–16.

4. Joung, J.K. and J.D. Sander, TALENs: a widely applicable technology for targeted genome editing. Nat Rev Mol Cell Biol, 2013. 14(1): p. 49–55.

5. Carroll, D., Genome engineering with zinc-finger nucleases. Genetics, 2011. 188(4): p. 773–82.

6. Hofer, U., J.E. Henley, C.M. Exline, O. Mulhern, E. Lopez and P.M. Cannon, Pre-clinical modeling of CCR5 knockout in human hematopoietic stem cells by zinc finger nucleases using humanized mice. J Infect Dis, 2013. 208 Suppl 2: p. S160–4.

7. Romito, M., A. Juillerat, Y.L. Kok, M. Hildenbeutel, M. Rhiel, G. Andrieux, J. Geiger, C. Rudolph, C. Mussolino, A. Duclert, K.J. Metzner, P. Duchateau, T. Cathomen and T.I. Cornu, Preclinical Evaluation of a Novel TALEN Targeting CCR5 Confirms Efficacy and Safety in Conferring Resistance to HIV-1 Infection. Biotechnol J, 2021. 16(1): p. e2000023.

8. Shi, B., J. Li, X. Shi, W. Jia, Y. Wen, X. Hu, F. Zhuang, J. Xi and L. Zhang, TALEN-Mediated Knockout of CCR5 Confers Protection Against Infection of Human Immunodeficiency Virus. J Acquir Immune Defic Syndr, 2017. 74(2): p. 229–241.

9. Tebas, P., D. Stein, W.W. Tang, I. Frank, S.Q. Wang, G. Lee, S.K. Spratt, R.T. Surosky, M.A. Giedlin, G. Nichol, M.C. Holmes, P.D. Gregory, D.G. Ando, M. Kalos, R.G. Collman, G. Binder-Scholl, G. Plesa, W.T. Hwang, B.L. Levine and C.H. June, Gene editing of CCR5 in autologous CD4 T cells of persons infected with HIV. N Engl J Med, 2014. 370(10): p. 901–10.

10. Cannon, P. and C. June, Chemokine receptor 5 knockout strategies. Curr Opin HIV AIDS, 2011. 6(1): p. 74–9.

11. Maier, D.A., A.L. Brennan, S. Jiang, G.K. Binder-Scholl, G. Lee, G. Plesa, Z. Zheng, J. Cotte, C. Carpenito, T. Wood, S.K. Spratt, D. Ando, P. Gregory, M.C. Holmes, E.E. Perez, J.L. Riley, R.G. Carroll, C.H. June and B.L. Levine, Efficient clinical scale gene modification via zinc finger nuclease-targeted disruption of the HIV co-receptor CCR5. Hum Gene Ther, 2013. 24(3): p. 245–58.

12. Knipping, F., G.A. Newby, C.R. Eide, A.N. McElroy, S.C. Nielsen, K. Smith, Y. Fang, T.I. Cornu, C. Costa, A. Gutierrez-Guerrero, S.P. Bingea, C.J. Feser, B. Steinbeck, K.L. Hippen, B.R. Blazar, A. McCaffrey, C. Mussolino, E. Verhoeyen, J. Tolar, D.R. Liu and M.J. Osborn, Disruption of HIV-1 co-receptors CCR5 and CXCR4 in primary human T cells and hematopoietic stem and progenitor cells using base editing. Mol Ther, 2022. 30(1): p. 130–144.

13. Huang, T.P., G.A. Newby and D.R. Liu, Precision genome editing using cytosine and adenine base editors in mammalian cells. Nat Protoc, 2021. 16(2): p. 1089–1128.

14. Jinek, M., K. Chylinski, I. Fonfara, M. Hauer, J.A. Doudna and E. Charpentier, A programmable dual-RNA-guided DNA endonuclease in adaptive bacterial immunity. Science, 2012. 337(6096): p. 816–21.

15. Gilbert, L.A., M.H. Larson, L. Morsut, Z. Liu, G.A. Brar, S.E. Torres, N. Stern-Ginossar, O. Brandman, E.H. Whitehead, J.A. Doudna, W.A. Lim, J.S. Weissman and L.S. Qi, CRISPR-mediated modular RNA-guided regulation of transcription in eukaryotes. Cell, 2013. 154(2): p. 442–51.

16. Cong, L., F.A. Ran, D. Cox, S. Lin, R. Barretto, N. Habib, P.D. Hsu, X. Wu, W. Jiang, L.A. Marraffini and F. Zhang, Multiplex genome engineering using CRISPR/Cas systems. Science, 2013. 339(6121): p. 819–23.

17. Dash, P.K., R. Kaminski, R. Bella, H. Su, S. Mathews, T.M. Ahooyi, C. Chen, P. Mancuso, R. Sariyer, P. Ferrante, M. Donadoni, J.A. Robinson, B. Sillman, Z. Lin, J.R. Hilaire, M. Banoub, M. Elango, N. Gautam, R.L. Mosley, L.Y. Poluektova, J. McMillan, A.N. Bade, S. Gorantla, I.K. Sariyer, T.H. Burdo, W.B. Young, S. Amini, J. Gordon, J.M. Jacobson, B. Edagwa, K. Khalili and H.E. Gendelman, Sequential LASER ART and CRISPR Treatments Eliminate HIV-1 in a Subset of Infected Humanized Mice. Nat Commun, 2019. 10(1): p. 2753.

18. Yang, Y.C. and H.C. Yang, Recent Progress and Future Prospective in HBV Cure by CRISPR/Cas. Viruses, 2021. 14(1).

19. Stone, D., K.R. Long, M.A. Loprieno, H.S. De Silva Feelixge, E.J. Kenkel, R.M. Liley, S. Rapp, P. Roychoudhury, T. Nguyen, L. Stensland, R. Colon-Thillet, L.M. Klouser, N.D. Weber, C. Le, J. Wagoner, E.A. Goecker, A.Z. Li, K. Eichholz, L. Corey, D.L. Tyrrell, A.L. Greninger, M.L. Huang, S.J. Polyak, M. Aubert, J.E. Sagartz and K.R. Jerome, CRISPR-Cas9 gene editing of hepatitis B virus in chronically infected humanized mice. Mol Ther Methods Clin Dev, 2021. 20: p. 258–275.

20. Lu, Y., J. Xue, T. Deng, X. Zhou, K. Yu, L. Deng, M. Huang, X. Yi, M. Liang, Y. Wang, H. Shen, R. Tong, W. Wang, L. Li, J. Song, J. Li, X. Su, Z. Ding, Y. Gong, J. Zhu, Y. Wang, B. Zou, Y. Zhang, Y. Li, L. Zhou, Y. Liu, M. Yu, Y. Wang, X. Zhang, L. Yin, X. Xia, Y. Zeng, Q. Zhou, B. Ying, C. Chen, Y. Wei, W. Li and T. Mok, Safety and feasibility of CRISPR-edited T cells in patients with refractory non-small-cell lung cancer. Nat Med, 2020. 26(5): p. 732–740.

21. Vazquez-Salat, N., N. Yuhki, T. Beck, S.J. O’Brien and W.J. Murphy, Gene conversion between mammalian CCR2 and CCR5 chemokine receptor genes: a potential mechanism for receptor dimerization. Genomics, 2007. 90(2): p. 213–24.

22. Xu, L., H. Yang, Y. Gao, Z. Chen, L. Xie, Y. Liu, Y. Liu, X. Wang, H. Li, W. Lai, Y. He, A. Yao, L. Ma, Y. Shao, B. Zhang, C. Wang, H. Chen and H. Deng, CRISPR/Cas9-Mediated CCR5 Ablation in Human Hematopoietic Stem/Progenitor Cells Confers HIV-1 Resistance In Vivo. Mol Ther, 2017. 25(8): p. 1782–1789.

23. Gupta, R.K., S. Abdul-Jawad, L.E. McCoy, H.P. Mok, D. Peppa, M. Salgado, J. Martinez-Picado, M. Nijhuis, A.M.J. Wensing, H. Lee, P. Grant, E. Nastouli, J. Lambert, M. Pace, F. Salasc, C. Monit, A.J. Innes, L. Muir, L. Waters, J. Frater, A.M.L. Lever, S.G. Edwards, I.H. Gabriel and E. Olavarria, HIV-1 remission following CCR5Delta32/Delta32 haematopoietic stem-cell transplantation. Nature, 2019. 568(7751): p. 244–248.

24. Hutter, G., D. Nowak, M. Mossner, S. Ganepola, A. Mussig, K. Allers, T. Schneider, J. Hofmann, C. Kucherer, O. Blau, I.W. Blau, W.K. Hofmann and E. Thiel, Long-term control of HIV by CCR5 Delta32/Delta32 stem-cell transplantation. N Engl J Med, 2009. 360(7): p. 692–8.

25. Ding, J., Y. Liu and Y. Lai, Knowledge From London and Berlin: Finding Threads to a Functional HIV Cure. Front Immunol, 2021. 12: p. 688747.

26. Gupta, R.K., D. Peppa, A.L. Hill, C. Galvez, M. Salgado, M. Pace, L.E. McCoy, S.A. Griffith, J. Thornhill, A. Alrubayyi, L.E.P. Huyveneers, E. Nastouli, P. Grant, S.G. Edwards, A.J. Innes, J. Frater, M. Nijhuis, A.M.J. Wensing, J. Martinez-Picado and E. Olavarria, Evidence for HIV-1 cure after CCR5Delta32/Delta32 allogeneic haemopoietic stem-cell transplantation 30 months post analytical treatment interruption: a case report. Lancet HIV, 2020. 7(5): p. e340–e347.

27. Mehta, A. and O.M. Merkel, Immunogenicity of Cas9 protein. J Pharm Sci, 2020. 109(1): p. 62–67.

28. Crudele, J.M. and J.S. Chamberlain, Cas9 immunity creates challenges for CRISPR gene editing therapies. Nat Commun, 2018. 9(1): p. 3497.

29. Charlesworth, C.T., P.S. Deshpande, D.P. Dever, J. Camarena, V.T. Lemgart, M.K. Cromer, C.A. Vakulskas, M.A. Collingwood, L. Zhang, N.M. Bode, M.A. Behlke, B. Dejene, B. Cieniewicz, R. Romano, B.J. Lesch, N. Gomez-Ospina, S. Mantri, M. Pavel-Dinu, K.I. Weinberg and M.H. Porteus, Identification of preexisting adaptive immunity to Cas9 proteins in humans. Nat Med, 2019. 25(2): p. 249–254.

30. Chen, J.S., E. Ma, L.B. Harrington, M. Da Costa, X. Tian, J.M. Palefsky and J.A. Doudna, CRISPR-Cas12a target binding unleashes indiscriminate single-stranded DNase activity. Science, 2018. 360(6387): p. 436–439.

31. Zetsche, B., J.S. Gootenberg, O.O. Abudayyeh, I.M. Slaymaker, K.S. Makarova, P. Essletzbichler, S.E. Volz, J. Joung, J. van der Oost, A. Regev, E.V. Koonin and F. Zhang, Cpf1 is a single RNA-guided endonuclease of a class 2 CRISPR-Cas system. Cell, 2015. 163(3): p. 759–71.

32. Strecker, J., S. Jones, B. Koopal, J. Schmid-Burgk, B. Zetsche, L. Gao, K.S. Makarova, E.V. Koonin and F. Zhang, Engineering of CRISPR-Cas12b for human genome editing. Nat Commun, 2019. 10(1): p. 212.

33. Shmakov, S., O.O. Abudayyeh, K.S. Makarova, Y.I. Wolf, J.S. Gootenberg, E. Semenova, L. Minakhin, J. Joung, S. Konermann, K. Severinov, F. Zhang and E.V. Koonin, Discovery and Functional Characterization of Diverse Class 2 CRISPR-Cas Systems. Mol Cell, 2015. 60(3): p. 385–97.

34. Chen, L.X., B. Al-Shayeb, R. Meheust, W.J. Li, J.A. Doudna and J.F. Banfield, Candidate Phyla Radiation Roizmanbacteria From Hot Springs Have Novel and Unexpectedly Abundant CRISPR-Cas Systems. Front Microbiol, 2019. 10: p. 928.

35. Burstein, D., L.B. Harrington, S.C. Strutt, A.J. Probst, K. Anantharaman, B.C. Thomas, J.A. Doudna and J.F. Banfield, New CRISPR-Cas systems from uncultivated microbes. Nature, 2017. 542(7640): p. 237–241.

36. Tsuchida, C.A., S. Zhang, M.S. Doost, Y. Zhao, J. Wang, E. O’Brien, H. Fang, C.P. Li, D. Li, Z.Y. Hai, J. Chuck, J. Brotzmann, A. Vartoumian, D. Burstein, X.W. Chen, E. Nogales, J.A. Doudna and J.G. Liu, Chimeric CRISPR-CasX enzymes and guide RNAs for improved genome editing activity. Mol Cell, 2022. 82(6): p. 1199–1209 e6.

37. Abudayyeh, O.O., J.S. Gootenberg, S. Konermann, J. Joung, I.M. Slaymaker, D.B. Cox, S. Shmakov, K.S. Makarova, E. Semenova, L. Minakhin, K. Severinov, A. Regev, E.S. Lander, E.V. Koonin and F. Zhang, C2c2 is a single-component programmable RNA-guided RNA-targeting CRISPR effector. Science, 2016. 353(6299): p. aaf5573.

38. Liu, J.J., N. Orlova, B.L. Oakes, E. Ma, H.B. Spinner, K.L.M. Baney, J. Chuck, D. Tan, G.J. Knott, L.B. Harrington, B. Al-Shayeb, A. Wagner, J. Brotzmann, B.T. Staahl, K.L. Taylor, J. Desmarais, E. Nogales and J.A. Doudna, CasX enzymes comprise a distinct family of RNA-guided genome editors. Nature, 2019. 566(7743): p. 218–223.

39. Liu, J.-J., N. Orlova, B.L. Oakes, E. Ma, H.B. Spinner, K.L.M. Baney, J. Chuck, D. Tan, G.J. Knott, L.B. Harrington, B. Al-Shayeb, A. Wagner, J. Brötzmann, B.T. Staahl, K.L. Talyor, J. Desmarais, E. Nogales and J.A. Doudna, CRISPR-CasX is an RNA-dominated enzyme active for human genome editing. Nature, 2019. 566(7743): p. 218–23.

40. Mummidi, S., S.S. Ahuja, B.L. McDaniel and S.K. Ahuja, The human CC chemokine receptor 5 (CCR5) gene. Multiple transcripts with 5’-end heterogeneity, dual promoter usage, and evidence for polymorphisms within the regulatory regions and noncoding exons. J Biol Chem, 1997. 272(49): p. 30662–71.

41. Liu, R., W.A. Paxton, S. Choe, D. Ceradini, S.R. Martin, R. Horuk, M.E. MacDonald, H. Stuhlmann, R.A. Koup and N.R. Landau, Homozygous defect in HIV-1 coreceptor accounts for resistance of some multiply-exposed individuals to HIV-1 infection. Cell, 1996. 86(3): p. 367–77.

42. Selkova, P., A. Vasileva, G. Pobegalov, O. Musharova, A. Arseniev, M. Kazalov, T. Zyubko, N. Shcheglova, T. Artamonova, M. Khodorkovskii, K. Severinov and I. Fedorova, Position of Deltaproteobacteria Cas12e nuclease cleavage sites depends on spacer length of guide RNA. RNA Biol, 2020. 17(10): p. 1472–1479.

43. Heredia, J.D., J. Park, R.J. Brubaker, S.K. Szymanski, K.S. Gill and E. Procko, Mapping Interaction Sites on Human Chemokine Receptors by Deep Mutational Scanning. J Immunol, 2018. 200(11): p. 3825–3839.

44. Hsu, P.D., D.A. Scott, J.A. Weinstein, F.A. Ran, S. Konermann, V. Agarwala, Y. Li, E.J. Fine, X. Wu, O. Shalem, T.J. Cradick, L.A. Marraffini, G. Bao and F. Zhang, DNA targeting specificity of RNA-guided Cas9 nucleases. Nat Biotechnol, 2013. 31(9): p. 827–32.

