## Supplemental Files for "CAS12e (CASX2) CLEAVAGE OF CCR5: IMPACT OF GUIDE RNA LENGTH AND PAM SEQUENCE ON CLEAVAGE ACTIVITY"

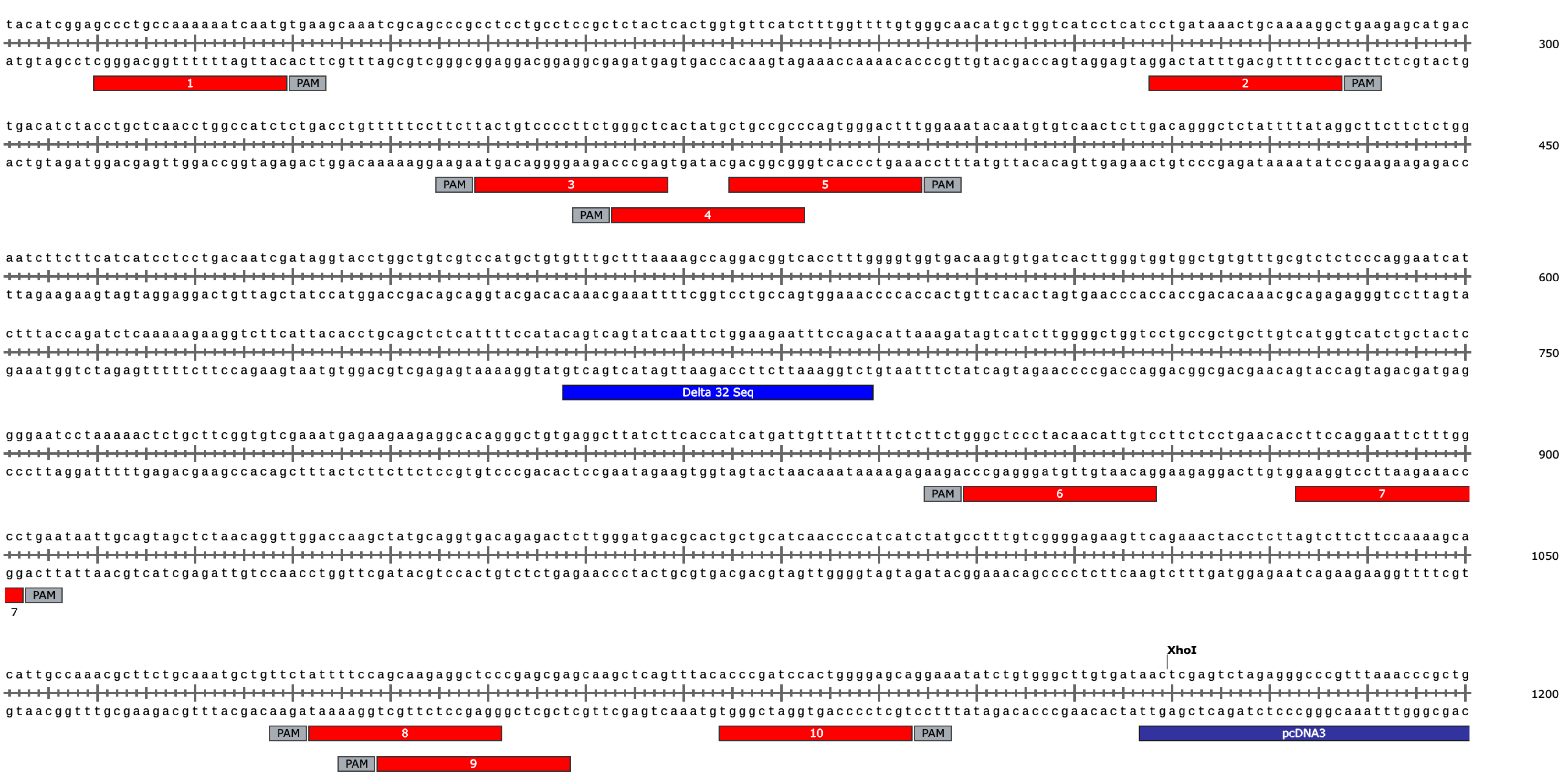


Supplemental Figure 1.

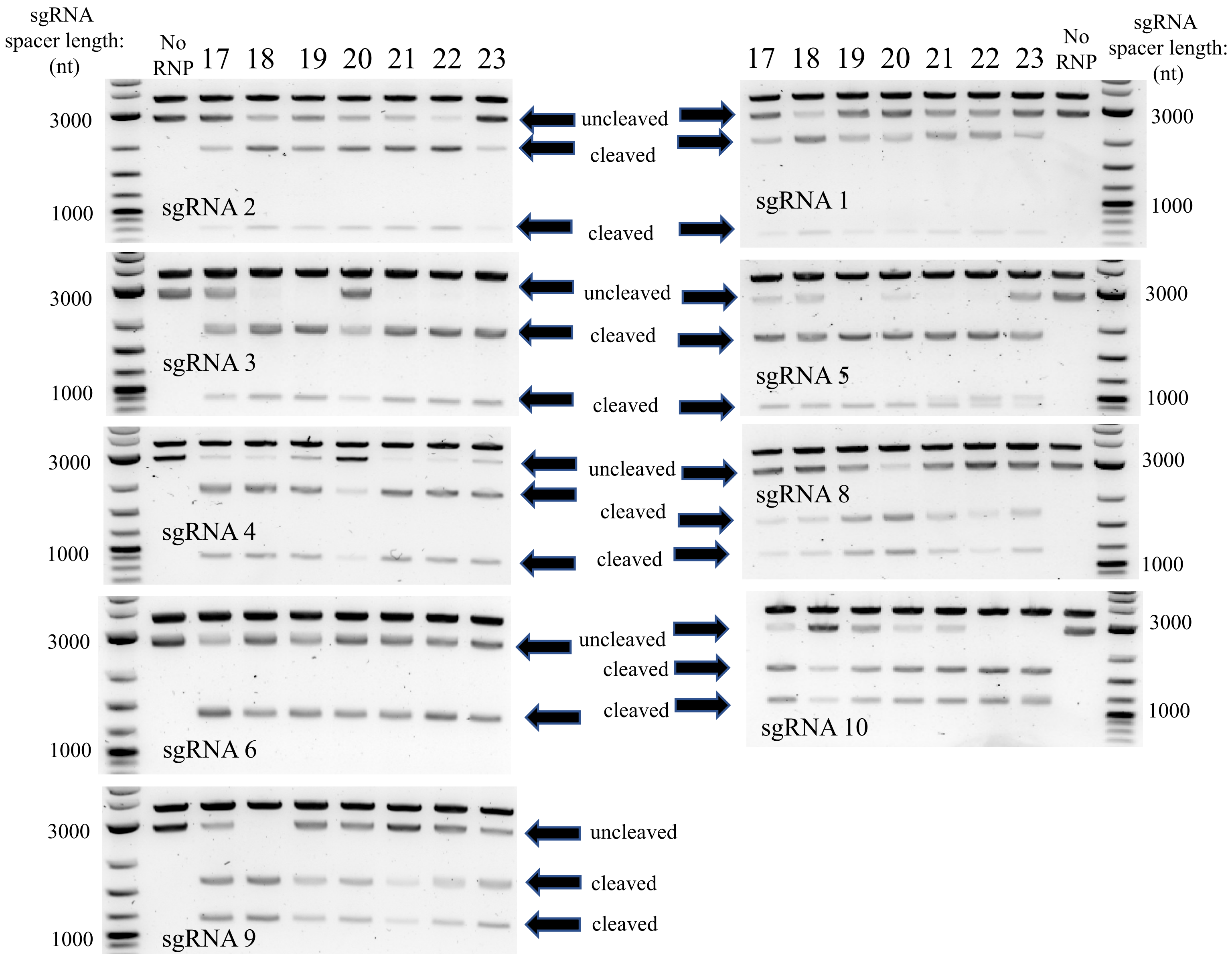


Supplemental Figure 2.

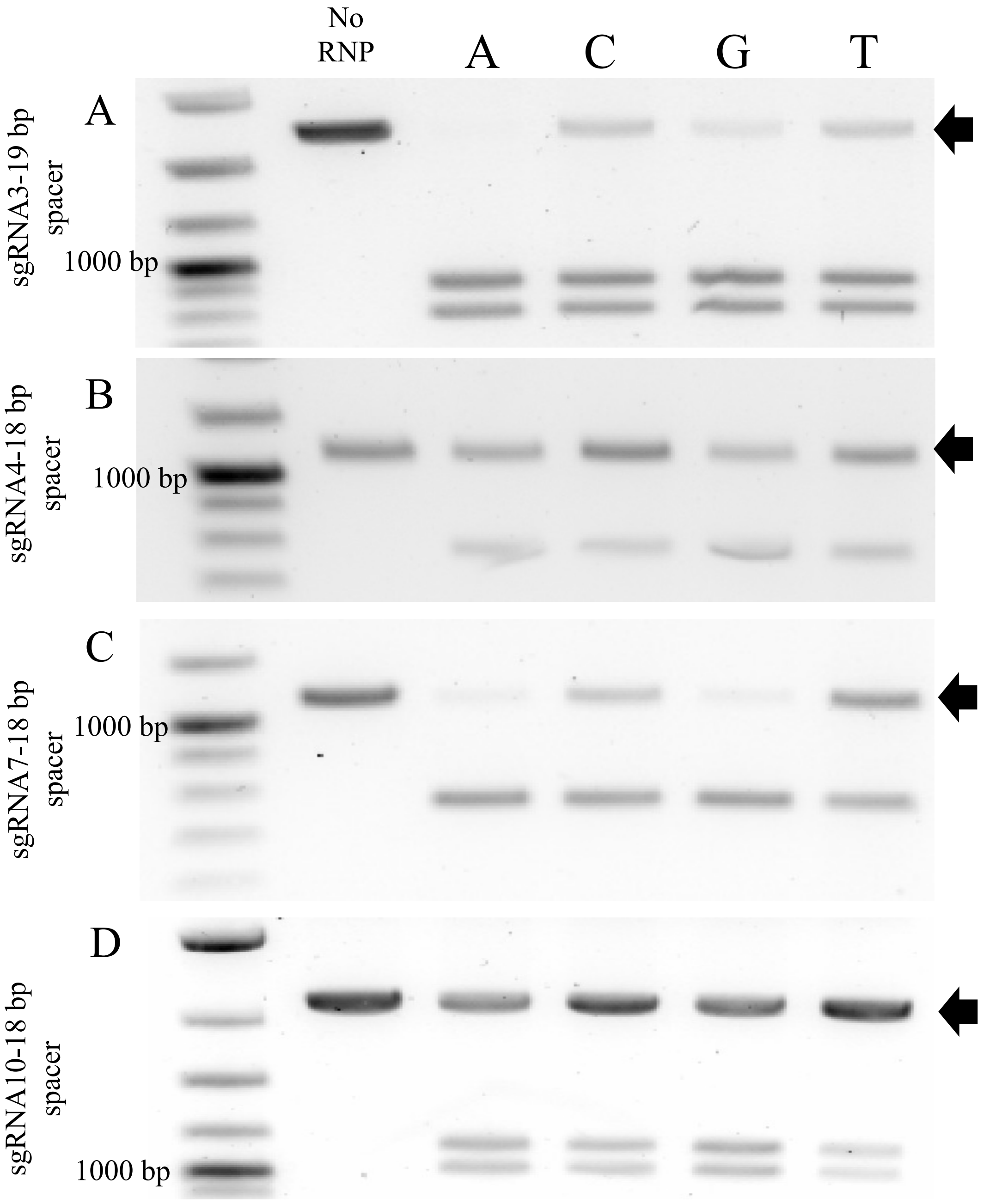


Supplemental Figure 3.

**Supplemental Table 1.** **DNA OLIGOS for in vitro transcription of guide RNAs**

### Scaffold DNA Oligo

5'- gaaatTAATACGACTCACTATAGTACTGGCGCTTTTATCTCATTACTTTGAGAGCCATCACCAGCGACTATGTCGTATGGGTAAAGCGCTTATTTATCGGAGAGAAATCC - 3'

TRACR RNA 5'-GTACTGGCGCTTTTATCTCATTACTTTGAGAGCCATCACCAGCGACTATGTCGTATGGGTAAAGCGCTTATTTATCGGAGA(81)

BLACK - T7 PRIMER

RED - TRACR

GREEN - LOOP/LINKER

### Reverse DNA Oligo

3'- ATAAATAGCCTCTCTTTAGGCTATTTATTCTTCGTAGTTTCNNNNNNNNNNNNNNNNNNNN-5'

PURPLE - (NNN's) - GUIDE REVERSE OLIGOS (17-23 BP)

BLUE - REPEAT

GREEN - LOOP/LINKER COMPLEMENT

**Supplemental Table 2. CCR5 sgRNA characteristics**

| Index | % Target Cleavage |
|---|---|
| (++++) | 90-100 |
| (+++) | 89-70 |
| (++) | 69-50 |
| (+) | <50 |

| sgRNA Number | Spacer Length | Guide Sequence | Terminal RNA Base | % G/C | PAM | Cutting Index | delta G (kcal/mol) |
|---|---|---|---|---|---|---|---|
| 1 | 17 | 5'-CAUUGAUUUUUUGGCAG | G | 35 | TTCA | (+) | -31.81 |
| | 18 | 5'-CAUUGAUUUUUUGGCAGG | G | 39 | | (++) | -34.88 |
| | 19 | 5'-CAUUGAUUUUUUGGCAGGG | G | 42 | | (+) | -37.95 |
| | 20 | 5'-CAUUGAUUUUUUGGCAGGGC | C | 45 | | (+) | -41.09 |
| | 21 | 5'-CAUUGAUUUUUUGGCAGGGCU | U | 43 | | (++) | -42.69 |
| | 22 | 5'-CAUUGAUUUUUUGGCAGGGCUC | C | 45 | | (++) | -44.26 |
| | 23 | 5'-CAUUGAUUUUUUGGCAGGGCUCC | C | 48 | | (+) | -47.33 |
| 2 | 17 | 5'-GCCUUUUGCAGUUUAUC | C | 41 | TTCA | (+) | -31.53 |
| | 18 | 5'-GCCUUUUGCAGUUUAUCA | A | 39 | | (++) | -33.48 |
| | 19 | 5'-GCCUUUUGCAGUUUAUCAG | G | 43 | | (+) | -35.08 |
| | 20 | 5'-GCCUUUUGCAGUUUAUCAGG | G | 45 | | (++) | -38.15 |
| | 21 | 5'-GCCUUUUGCAGUUUAUCAGGA | A | 43 | | (++) | -39.72 |
| | 22 | 5'-GCCUUUUGCAGUUUAUCAGGAU | U | 41 | | (+++) | -41.2 |
| | 23 | 5'-GCCUUUUGCAGUUUAUCAGGAUG | G | 43 | | (+) | -43.15 |
| 3 | 17 | 5'-UACUGUCCCCUUCUGGG | G | 59 | TTCT | (+++) | -32.79 |
| | 18 | 5'-UACUGUCCCCUUCUGGGC | C | 61 | | (++) | -35.93 |
| | 19 | 5'-UACUGUCCCCUUCUGGGCU | U | 58 | | (+++) | -37.53 |
| | 20 | 5'-UACUGUCCCCUUCUGGGCUC | C | 60 | | (+++) | -39.1 |
| | 21 | 5'-UACUGUCCCCUUCUGGGCUCA | A | 57 | | (++++) | -41.06 |
| | 22 | 5'-UACUGUCCCCUUCUGGGCUCAC | C | 59 | | (++++) | -42.4 |
| | 23 | 5'-UACUGUCCCCUUCUGGGCUCACU | U | 57 | | (++) | -44 |
| 4 | 17 | 5'-AAAGUCCCACUGGGCGG | G | 65 | TTCC | (+++) | -37.35 |
| | 18 | 5'-AAAGUCCCACUGGGCGGC | C | 67 | | (++++) | -40.49 |
| | 19 | 5'-AAAGUCCCACUGGGCGGCA | A | 63 | | (+++) | -42.44 |
| | 20 | 5'-AAAGUCCCACUGGGCGGCAG | G | 65 | | (+) | -44.04 |
| | 21 | 5'-AAAGUCCCACUGGGCGGCAGC | C | 67 | | (+++) | -47.18 |
| | 22 | 5'-AAAGUCCCACUGGGCGGCAGCA | A | 64 | | (+++) | -49.13 |
| | 23 | 5'-AAAGUCCCACUGGGCGGCAGCAU | U | 61 | | (+++) | -50.61 |
| 5 | 17 | 5'-GGGCUCACUAUGCUGCC | C | 65 | TTCT | (+++) | -34.64 |
| | 18 | 5'-GGGCUCACUAUGCUGCCG | G | 67 | | (++) | -38.25 |
| | 19 | 5'-GGGCUCACUAUGCUGCCGC | C | 68 | | (+++) | -41.39 |
| | 20 | 5'-GGGCUCACUAUGCUGCCGCC | C | 70 | | (+++) | -44.46 |
| | 21 | 5'-GGGCUCACUAUGCUGCCGCCC | C | 71 | | (++++) | -47.52 |
| | 22 | 5'-GGGCUCACUAUGCUGCCGCCCA | A | 68 | | (++++) | -49.48 |
| | 23 | 5'-GGGCUCACUAUGCUGCCGCCCAG | G | 70 | | (++) | -51.08 |

| | | | | | | | |
|---|---|---|---|---|---|---|---|
| 6 | 17 | 5'-GGGCUCCCUACAACAUU | U | 53 | TTCT | (+) | -33.1 |
| | 18 | 5'-GGGCUCCCUACAACAUUG | G | 56 | | (+) | -35.06 |
| | 19 | 5'-GGGCUCCCUACAACAUUGU | U | 53 | | (+++) | -36.4 |
| | 20 | 5'-GGGCUCCCUACAACAUUGUC | C | 55 | | (++) | -37.98 |
| | 21 | 5'-GGGCUCCCUACAACAUUGUCC | C | 57 | | (+) | -41.04 |
| | 22 | 5'-GGGCUCCCUACAACAUUGUCCU | U | 55 | | (+) | -42.64 |
| | 23 | 5'-GGGCUCCCUACAACAUUGUCCUU | U | 52 | | (+) | -44.59 |
| 7 | 17 | 5'-GCCCAAAGAAUUCCUGG | G | 53 | TTCA | (++++) | -34.92 |
| | 18 | 5'-GGCCAAAGAAUUCCUGGA | A | 50 | | (++++) | -36.5 |
| | 19 | 5'-GGCCAAAGAAUUCCUGGAA | A | 47 | | (++++) | -38.44 |
| | 20 | 5'-GGCCAAAGAAUUCCUGGAAG | G | 50 | | (++++) | -40.04 |
| | 21 | 5'-GGCCAAAGAAUUCCUGGAAGG | G | 52 | | (++) | -43.11 |
| | 22 | 5'-GGCCAAAGAAUUCCUGGAAGGU | U | 50 | | (++) | -44.45 |
| | 23 | 5'-GGCCAAAGAAUUCCUGGAAGGUG | G | 52 | | (++) | -46.41 |
| 8 | 17 | 5'-AUUUUCCAGCAAGAGGC | C | 47 | TTCT | (+) | -33.52 |
| | 18 | 5'-AUUUUCCAGCAAGAGGCU | U | 44 | | (+) | -35.12 |
| | 19 | 5'-AUUUUCCAGCAAGAGGCUC | C | 47 | | (++) | -36.7 |
| | 20 | 5'-AUUUUCCAGCAAGAGGCUCC | C | 50 | | (+++) | -39.76 |
| | 21 | 5'-AUUUUCCAGCAAGAGGCUCCC | C | 52 | | (+) | -42.83 |
| | 22 | 5'-AUUUUCCAGCAAGAGGCUCCCG | G | 55 | | (+) | -46.45 |
| | 23 | 5'-AUUUUCCAGCAAGAGGCUCCCGA | A | 52 | | (+) | -48.02 |
| 9 | 17 | 5'-AGCAAGAGGCUCCCGAG | G | 65 | TTCC | (++) | -35.71 |
| | 18 | 5'-AGCAAGAGGCUCCCGAGC | C | 67 | | (++++) | -38.85 |
| | 19 | 5'-AGCAAGAGGCUCCCGAGCG | G | 68 | | (+) | -42.46 |
| | 20 | 5'-AGCAAGAGGCUCCCGAGCGA | A | 65 | | (++) | -44.04 |
| | 21 | 5'-AGCAAGAGGCUCCCGAGCGAG | G | 67 | | (+) | -45.64 |
| | 22 | 5'-AGCAAGAGGCUCCCGAGCGAGC | C | 68 | | (++) | -48.78 |
| | 23 | 5'-AGCAAGAGGCUCCCGAGCGAGCA | A | 65 | | (++) | -50.73 |
| 10 | 17 | 5'-UGCUCCCCAGUGGAUCG | G | 65 | TTCC | (+++) | -35.63 |
| | 18 | 5'-UGCUCCCCAGUGGAUCGG | G | 67 | | (+) | -38.7 |
| | 19 | 5'-UGCUCCCCAGUGGAUCGGG | G | 68 | | (++) | -41.77 |
| | 20 | 5'-UGCUCCCCAGUGGAUCGGGU | U | 65 | | (+++) | -43.11 |
| | 21 | 5'-UGCUCCCCAGUGGAUCGGGUG | G | 67 | | (+++) | -45.06 |
| | 22 | 5'-UGCUCCCCAGUGGAUCGGGUGU | U | 64 | | (++++) | -46.4 |
| | 23 | 5'-UGCUCCCCAGUGGAUCGGGUGUA | A | 61 | | (++++) | -47.36 |
